# When higher carrying capacities lead to faster propagation

**DOI:** 10.1101/307322

**Authors:** Marjorie Haond, Thibaut Morel-Journel, Eric Lombaert, Elodie Vercken, Ludovic Mailleret, Lionel Roques

**Affiliations:** Université Côte d’Azur, INRA, CNRS, ISA, France; Earth and Life Institute, Biodiversity Research Centre, Université Catholique de Louvain, Louvain-la-Neuve, Belgium; Inria, Biocore, 06902 Sophia Antipolis, France; BioSP, INRA, 84000, Avignon, France

**Keywords:** expansion dynamics, invasion biology, propagation speed, density-dependent dispersal, Allee effect, demographic stochasticity, experimental microcosms, small populations, pushed invasion

## Abstract

This preprint has been reviewed and recommended by Peer Community In Ecology (https://dx.doi.org/10.24072/pci.ecology.100004). Finding general patterns in the expansion of natural populations is a major challenge in ecology and invasion biology. Classical spatio-temporal models predict that the carrying capacity (*K*) of the environment should have no influence on the speed (*v*) of an expanding population. We tested the generality of this statement with reaction-diffusion equations, stochastic individual-based models, and microcosms experiments with *Trichogramma chilonis* wasps. We investigated the dependence between *K* and *v* under different assumptions: null model (Fisher-KPP-like assumptions), strong Allee effects, and positive density-dependent dispersal. These approaches led to similar and complementary results. Strong Allee effects, positive density-dependent dispersal and demographic stochasticity in small populations lead to a positive dependence between *K* and *v*. A positive correlation between carrying capacity and propagation speed might be more frequent than previously expected, and be the rule when individuals at the edge of a population range are not able to fully drive the expansion.

## 1. Introduction

A significant acceleration in the rate of introduction of alien species has been observed since the 19th century (Seebens et al., 2017). After their establishment, invading species spread by expanding their range into suitable environment (Blackburn et al., 2011), with sometimes important impacts on agriculture production, biodiversity or human health (Crooks, 2002; Keller et al., 2011). The need for accurate predictions in order to anticipate range shifts and manage populations is ever increasing (Hulme, 2009; Schwartz, 2012). Yet this aim has proven difficult to achieve, because propagation speed and expansion patterns are highly variable across populations and depend on many different factors (Hastings et al., 2005; Hui et al., 2011).

Studies on introduced species often show that invasion success is correlated with life-history traits (Moravcova et al., 2010; van Kleunen et al., 2011; Gidoin et al., 2015) yet, the properties of the environment also affect expansion patterns. Previous studies have analyzed the impact of habitat quality on expansion. They focused on the variations in the population growth rate across space as a proxy for habitat quality (Shigesada and Kawasaki, 1997; Neubert and Caswell, 2000; Kanarek et al., 2008; Mortelliti et al., 2010). The carrying capacity is another component of habitat quality, thought to be deeply impacting establishment dynamics (Drake and Lodge, 2006; Verbruggen et al., 2013; Vercken et al., 2013) and extinction probability (Griffen and Drake, 2008; Belovsky et al., 1999). However carrying capacity has not been investigated in connection with expansion speed so far, probably because standard theoretical models predict no relationship between these two quantities.

Reaction-diffusion models are recognized as robust descriptors of the qualitative properties of many ecological systems (e.g. Turchin, 1998; Hastings et al., 2005; Gilad et al., 2007). In such models, the carrying capacity, usually denoted *K*, is a positive stable equilibrium corresponding to the maximum population density that can be sustained by the environment at any location *x*. One of the most classical model is the “Fisher-KPP” model, with a logistic growth *f*(*u*) = *r u*(1 – *u/K*) (*u* being the population density at time *t* and location *x*, and *r* the intrinsic growth rate). It has been widely used to describe the spatio-temporal dynamics of expanding populations (e.g. Shigesada and Kawasaki, 1997, and the references therein). In this model, it is well-known since the pioneering work of Kolmogorov et al. (1937) that the speed of range expansion only depends on the growth function *f* through the limit of the per capita population growth rate *f*(*u*)*/u* as *u →* 0, i.e., through *f ′*(0). This means that the growth of populations with intermediate densities (*u > ε >* 0) has no effect on the propagation speed. Thus, expansion is driven by individuals at small density at the edge of the population range, where intraspecific competition vanishes. A direct consequence is that the speed of range expansion does not depend on the carrying capacity *K*. It is fully determined by the intrinsic growth rate and the diffusion coefficient leading to simple formulas. These formulas have been used to calculate theoretical expansion speeds for historical datasets on invasive populations (e.g. Van den Bosch et al., 1992; Holmes, 1993). Yet several empirical observations were not consistent with such predictions, in particular in presence of demographic properties like the Allee effect (Okubo et al., 1989; Liebhold et al., 1992; Neubert and Caswell, 2000). In such cases, it appeared that population propagation was not entirely driven by individuals at the edge of the range. Thus these propagations could not be described satisfactorily by the Fisher-KPP model.

There are two main reasons why these individuals at the edge of a population range might not be able to fully drive the expansion. Firstly, those individuals do not disperse (Altwegg et al., 2013), or secondly, their growth rate is limited, e.g. due to unfavorable conditions (Owen and Lewis, 2001; Silva et al., 2002; Garnier and Lewis, 2016), limited reproduction (Austerlitz et al., 2000; Courchamp et al., 2008) or competition (Roques et al., 2015). In such situations, individuals from high-density areas are involved in the expansion. We could therefore expect a positive relationship between carrying capacity and propagation speed. We investigate this hypothesis by analyzing propagation dynamics in presence of three factors known to penalize colonization success in small populations: strong Allee effects, positive density-dependent dispersal and demographic stochasticity.

A strong Allee effect induces a negative growth rate for densities lower than some value called the Allee threshold. This is widely studied, and observed in some animals as well as in some plants (Courchamp et al., 2008). It has been associated with reduced spread rates in invasive populations (Veit and Lewis, 1996; Davis et al., 2004; Taylor and Hastings, 2005). Positive density-dependent dispersal consists in an increase of the individual probability to disperse when the population density gets larger. Typically this kind of dispersal is supposed to allow organisms to avoid intraspecific competition or sexual harassment for females (Matthysen, 2005). This is common in mammals, birds and insects and may slow down rates of range expansion (Travis et al., 2009). Demographic stochasticity is likely to affect all small populations without any specific ecological mechanism. When local population size is small, the probability that no individual manages to successfully disperse and reproduce beyond the population range is increased, leading to a reduced propagation speed (Brunet and Derrida, 1997; Snyder, 2003).

The main objective of this work is to assess whether each of these three factors leads to a dependence of the population propagation speed *v* on *K*. In order to give more robustness to our study, we base our answers on two complementary modeling frameworks and an experimental approach. The first modeling framework is based on reaction-diffusion equations, which have the advantage of leading to simple formulas connecting *K* to *v*. The second modeling framework is based on stochastic individual-based simulation models (IBMs). Though less analytically tractable, these models are often considered as more realistic when dealing with small population sizes. The experimental study is made on a parasitoid wasp in laboratory microcosms, for which we establish that positive density-dependent dispersal occurs.

## 2. Material and Method

The next three sections are dedicated to the presentation of the two modeling frameworks and of the experimental study that we use to analyze the dependence between the carrying capacity and the propagation speed of a range-expanding population. For each modeling framework, we begin with a general presentation of the models, followed by a description of the way the three main scenarios (null model, strong Allee effect, positive density-dependent dispersal) are modeled.

### 2.1 Reaction-diffusion models

In one-dimensional reaction-diffusion models, the population density at time *t* and spatial location *x* is described by a function *u*(*t, x*) which satisfies a partial differential equation:

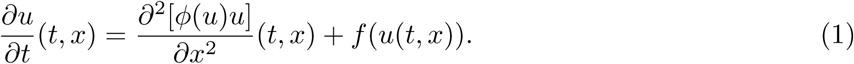

The operator *∂*^2^*/∂x*^2^ is the 1-dimensional Laplace diffusion operator. It describes uncorrelated random walk movements of the individuals, whose mobility is measured by the diffusion coefficient *ϕ*(*u*), which may depend on *u* or not, depending on the presence of density-dependent dispersal. The function *f* describes local population growth. The state 0 is where the species is not present, and the state *K* (carrying capacity) is where the population does not grow locally: *f* (0) = *f* (*K*) = 0.

The asymptotic propagation speed (or propagation speed, for short) to the right is the only speed *v* such that any observer who travels to the right – or to the left – with a speed larger than *v* will eventually see the population density go to 0, whereas any observer traveling with a speed slower than *v* will eventually see the density approach the carrying capacity *K*. Under the assumptions that are detailed below, when *ϕ*(*u*) = *D* is constant, it is known that the propagation speed exists and is finite (Kolmogorov et al., 1937; Aronson and Weinberger, 1975; Fife and McLeod, 1977).

#### Null model: Fisher-KPP

We assume a constant diffusion coefficient *ϕ*(*u*) = *D* (density-independent dispersal) and a growth function *f* such that the per capita growth rate *f* (*u*)*/u* reaches its maximum *r >* 0 when *u* approaches 0: 0 *< f* (*u*) *≤ ru* for all *u ∈* (0, *K*), with *r* = *f′*(0). This means that there is no Allee effect. In this case, the propagation speed is (Kolmogorov et al., 1937):

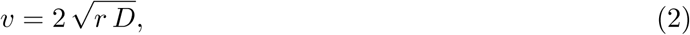

and therefore does not depend on *K*. A typical example is the logistic growth function *f* (*u*) =*r u*(1 – *u/K*).

#### Strong Allee effect

Again, we assume a constant diffusion coefficient *ϕ*(*u*) = *D*. Strong Allee effects are modeled by growth functions *f* (*u*) that are negative when *u* is below some threshold *ρ >* 0 (Allee threshold), and positive when *u* is between *ρ* and *K*. In this case, propagation can only occur (i.e., *v >* 0) when the average value of *f* over (0, *K*) is positive 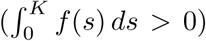. We believe that the standard form of the function *f* used e.g. by Lewis and Kareiva (1993) and Turchin (1998) is not useful for the purpose of this study. With this functional form,

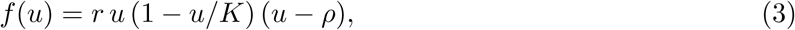

with *K > ρ >* 0, the maximum per capita growth rate max 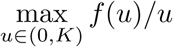 is equal to *r*(*K – ρ*)^2^*/*(4*K*). Thus, increasing *K* also induces a linear increase in the maximum per capita growth rate. In order to disentangle the effect of the carrying capacity from the effect of the per capita growth rate, we propose a new form for the growth function *f* :

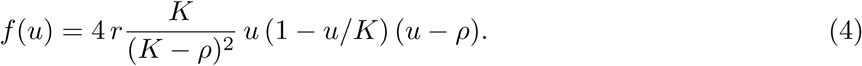

With this function *f*, the maximum sustainable density is still equal to *K*. However, contrarily to what we observed with the form (3), the maximum per capita growth rate is independent of *K* and equal to *r*. The average per capita growth rate in the interval *u ∈* (*ρ, K*) is also independent of *K*, and equal to 2 *r/*3. Thus, the new functional form (4) allows us to study the effect of the carrying capacity per se, i.e., when K has a limited effect on the growth of populations under optimal density conditions. A graphical comparison of the standard form of *f* (3) and the new form (4) is provided in Supplementary Information S1.

#### Positive density-dependent dispersal

We assume a simple logistic growth function *f* (*u*) = *r u*(1 – *u/K*) (no Allee effect). Positive density-dependent dispersal is modeled by considering an increasing function *ϕ*(*u*). The most standard form is *ϕ*(*u*) = *D u*^*a*^, with *D, a >* 0, the corresponding reaction-diffusion equation being the “porous media equation” (Vázquez, 2007). We focus here on the case *a* = 1, which was in part analyzed by Murray (2002), and for which an explicit formula for the speed is available in the particular case *K* = 1 (Newman, 1980), and can easily be adapted to any *K >* 0.

Other forms of *ϕ*(*u*) could be considered as well. For example *ϕ*(*u*) = *u*^*a*^*/*(*τ*^*a*^ + *u*^*a*^) with *τ, a >* 0 may be an appropriate function to describe a saturation effect (*ϕ*(*u*) *→* 1 for large values of *u*) but there is no general formula for the propagation speed in this case.

### 2.2 Individual-based stochastic simulation models (IBMs)

We consider a discrete time and discrete space stepping-stone model on a one-dimensional infinite grid indexed by *i ∈ ℕ*: the focus is on the propagation to the right, as in the theoretical models of Section 2.1. The number of individuals on the patch *i* of the grid, at time *t* is denoted by *N*_*i*_(*t*). We assume non-overlapping generations (of duration *δt* = 1). The population distribution at time *t* + 1 is obtained from three consecutive steps: reproduction, dispersal and competition, described below.

*Reproduction step.* The number of offspring at each position *i ≥* 0 is a random variable following a Poisson distribution with mean *R g*(*N*_*i*_(*t*)):

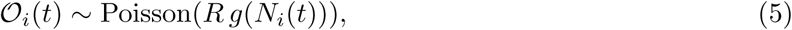

for a function *g* which depends on the assumptions (Allee effect or not) and with *R* the mean number of offspring per individual, per generation, in optimal conditions. Those *Dispersal step.* We assume random walk movements of individuals. Since generations are non-overlapping, only movements of the offspring are considered. Thus, at each generation, each individual stays at the same position with probability *p*_0_, or migrates to an adjacent patch with probability *p*_1_ = 1 – *p*_0_ (same probability *p*_1_/2 to move to the left or to the right), which may depend on the local population size *𝒪*_*i*_ or not, depending on the assumptions (presence of density-dependent dispersal or not). At each position *i*, the numbers of offspring moving to the left 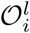, staying at the same position 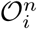, and moving to the right 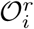 follow a multinomial distribution 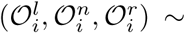, and the number of individuals *𝒟*_*i*_ at position *i* after the dispersion is equal to 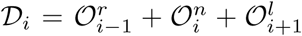. Note that, for small time steps *δ*_*t*_ and space steps *δ*_*x*_ (here, *δ*_*t*_ = *δ*_*x*_ = 1), this type of dispersal can be approached by a diffusion operator, of the same form as in Section 2.1, with e.g, 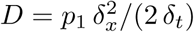 when *p*_1_ is constant (see Roques, 2013, Chapter 2 and references therein).

*Competition step.* As our objective is to understand the effect of the carrying capacity per se, we assume that *K* has no impact on the reproduction step. Thus, the competition step is the only step which is influenced by the carrying capacity *K*. To avoid an effect of *K* on the maximum per capita growth rate, we consider an extreme case where the effect of competition is negligible when the population size is below the threshold *K* and then increases continuously.

By definition of the carrying capacity, the expected number of deaths due to competition should be equal to the number of individuals exceeding *K*, at each position. The number of deaths due to competition *µ*_*i*_ is 0 if 𝒟_*i*_ *< K*, and follows a binomial distribution otherwise: *µ*_*i*_ *∼* Binomial(𝒟_*i*_, 1 – *K/𝒟*_*i*_). This implies that the expected number of deaths is *E*[*µ*_*i*_*|𝒟*_*i*_] = 𝒟_*i*_ – *K* when 𝒟_*i*_ *≥ K*. Then, the population distribution at generation *t* + 1 can be computed: *N*_*i*_(*t* + 1) = max(𝒟_*i*_ – *µ*_*i*_, 0).

We now describe the reproduction and dispersal steps under the three considered scenarios.

#### Null model: density-independent dispersal, no Allee effect

In the reproduction step, we simply assume that *g*(*N*_*i*_) = *N*_*i*_. With this assumption, the expected per capita offspring number *E*(𝒪_*i*_*/N*_*i*_) is constant equal to *R* (Fig. 1). In the dispersal step, we assume that *p*_0_ (and therefore *p*_1_) are independent of the local population size.

**Figure 1:**
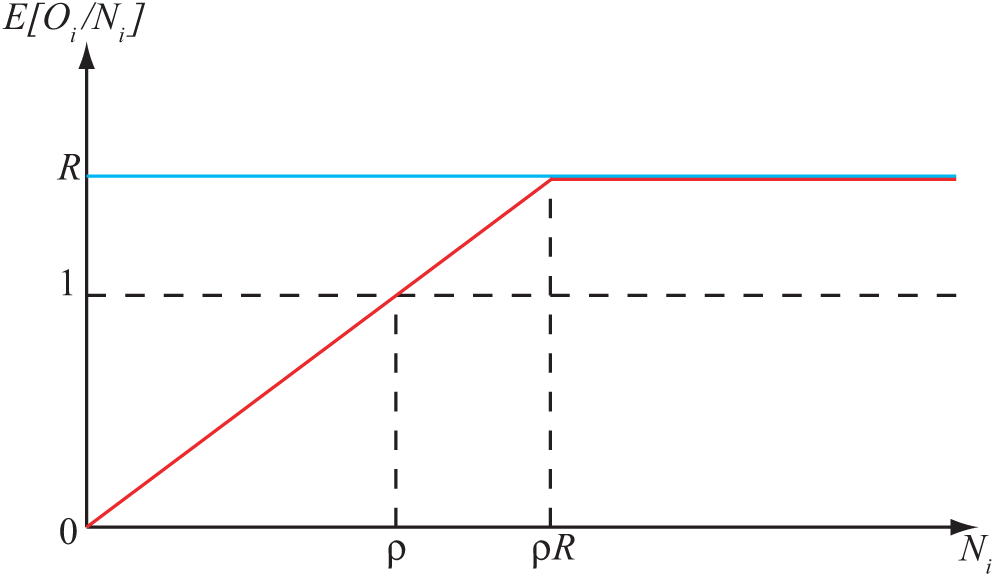
Schematic plot of the expected per capita offspring *E*(𝒪_*i*_*/N*_*i*_) in terms of *N*_*i*_. Horizontal blue line: no Allee effect. Red line: Allee effect with threshold *ρ*. *R* is the average per capita growth rate.

#### Strong Allee effect

In the reproduction step, we assume

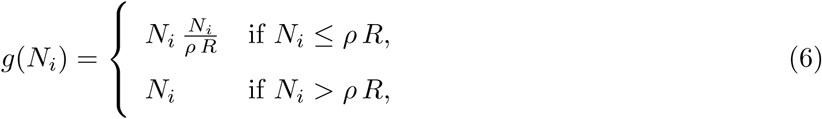

for for an integer *ρ ≥* 2. The corresponding shapes of the expected per capita offspring *E*(𝒪_*i*_*/N*_*i*_) are depicted in Fig. 1. The expected offspring number is larger than the parent population if and only if *N*_*i*_ is larger than the threshold *ρ*. Note that, with these assumptions, the maximum expected per capita offspring number in the IBM is equal to *R*, and is therefore independent of *K*. This is consistent with the reaction-diffusion framework (4).

A value *ρ* = 1 would correspond to a weak Allee effect: due to the discrete nature of *N*_*i*_, the per capita growth rate is always larger than 1, but the maximum is reached for some *N*_*i*_ = *ρ R >* 1.

#### Positive density-dependent dispersal

In this scenario, we assume that there is no Allee effect (*g*(*N*_*i*_) = *N*_*i*_) and that *p*_1_ is an increasing function of the number of individuals 𝒪_*i*_, following a sigmoid-like dependence:

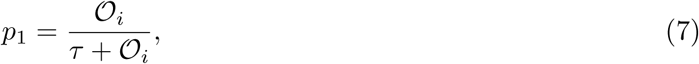

for some constant *τ* that will be specified later. Thus, 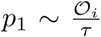 for small values of 𝒪_*i*_ and *p*_1_ = 1/2 when 𝒪_*i*_ = *τ* and the probability *p*_0_ = 1 – *p*_1_ is well-defined: *p*_0_ *∈* [0, 1].

In all cases, we assumed a boundary condition *N*_0_(*t*) = *K* and initial condition *N*_*i*_(0) = 0 for *i ≥* 1. The speed was computed as the number of colonized patches (i.e., with population larger than *K/*10) over the number of generations (300 generations). Because the dispersal is local (with *δ*_*x*_ = 1), the speed cannot exceed 1: at most one new patch is colonized per generation. In each scenario, for each parameter value (to be specified later, in the Results Section), and for any *K* ∈ {1, … , 500}, we carried out 200 replicate simulations, and measured the mean speed 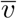 among these replicates together with 99% confidence intervals for the mean. In all cases, we fixed *R* = *e*^1^, corresponding to the analogue of *r* = 1 in the continuous time approaches of Section 2.1. Examples of simulated range expansions are given in Fig. 2.

**Figure 2:**
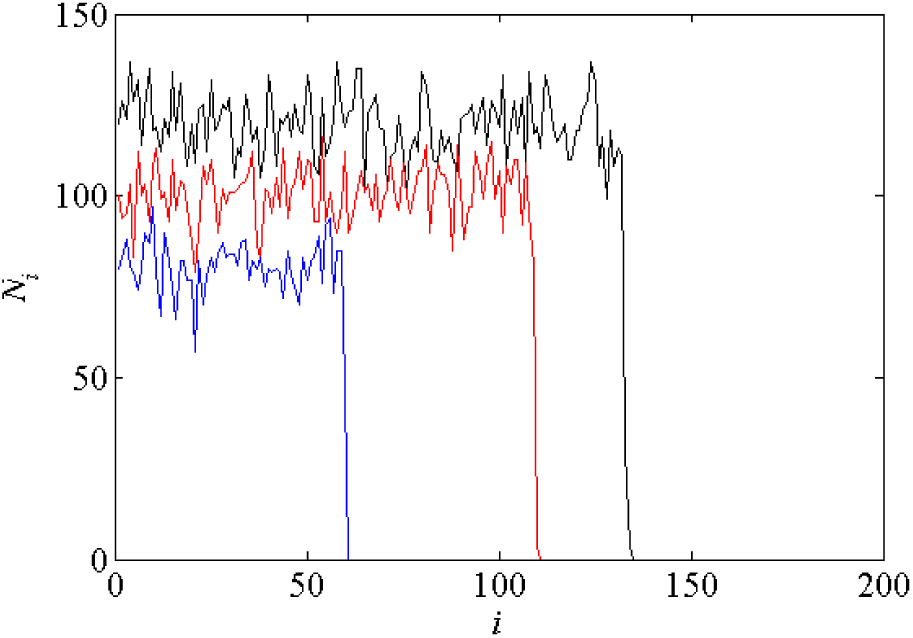
Range expansions simulated with the IBMs. Black curve: null model, with *K* = 120 and probability of migration *p*_1_ = 0.5 ; red curve: strong Allee effect, *ρ* = 20 and *K* = 100, *p*_1_ = 0.5; blue curve: density-dependent dispersal, *τ* = 500, *K* = 80, *p*_1_ = 0.14. In all cases, *t* = 150.

### 2.3 Experimental study: propagation of a parasitoid wasp in a microcosm

To test in experimental conditions whether positive density-dependent dispersal leads to a positive correlation between *K* and *v*, we carried out a microcosm experiment on the minute wasp *Trichogramma chilonis*, for which positive density-dependent dispersal has been documented together with the absence of Allee effect (Morel-Journel et al., 2016).

#### Biological model

We used a wasp from the Trichogrammatidae family. This hymenoptera of size less than 1mm is an egg parasitoid of several lepidopteran species. Trichogramma are used as biological control agents against different pests (Smith, 1996). They are well-suited for microcosm experiments because generation time is short and breeding is easy. In addition, parasitized eggs can be identified by their color which turns from white to dark gray because of chitinisation of the parasitoid pupa (Reay-Jones et al., 2006). As females lay at most one offspring per host eggs the number of parasitized eggs can be directly counted to estimate population size at the next generation.

#### Experimental protocol

The protocol is similar to the one established in Morel-Journel et al. (2016) with the same biological model. The experiment was conducted in climatic rooms with controlled temperature, lighting and humidity. Day cycles lasted 16 hours at 25*°*C, night cycles lasted 8 hours at 20*°*C, with a constant humidity around 70%. Populations of Trichogramma were introduced in experimental landscapes composed of 11 patches (see-through plastic tubes: high: 100mm, diameter: 50mm). Patches are connected by see-through plastic pipes (length: 400mm, diameter: 5mm) in order to create a one-dimensional stepping stone landscape. Initial populations were composed of 50 parasitized host eggs placed in the central patch of the landscape. Colonization occurred on both sides of this patch, see Fig. 3.

**Figure 3:**
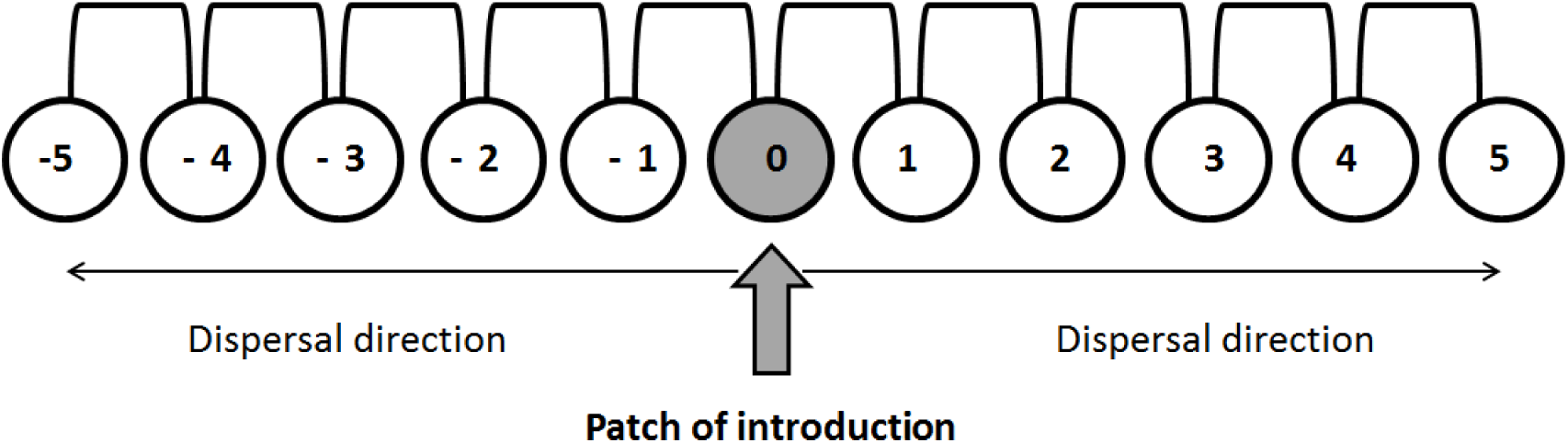
Experimental system: 11 plastic tubes connected with pipes to form a one dimensional stepping-stone landscape. Population can spread on both sides of the introduction patch. The propagation speed is computed as the number of colonized patches on one side divided by the number of generation.

Hosts were eggs of the Mediterranean flour moth, *Ephestia kuehniella*. The eggs were irradiated to prevent larval development of the moth while allowing the development of the trichogramma. Each generation cycle was completed in 9 days. The first step of the cycle was the emergence of adults. At this stage fresh host eggs were introduced, and each patch was connected to its direct neighbors to allow parasitoid migration. For the next two days, individuals could mate, lay eggs and possibly migrate to neighboring patches. At the end of those two days adults were removed in order to obtain non-overlapping generations. Host eggs exposed to parasitism were set aside until the emergence of the next generation of parasitoids. After four days, photographs of host eggs were taken, and parasitized eggs which had turned black at this stage, were counted with the help of the software ImageJ. The available number of host eggs in each patch determines the maximum parasitoid density in the next generation. Thus, it is directly correlated to the carrying capacity in this patch. In order to analyze the impact of the carrying capacity on propagation speed, we compared two modalities in the number of host eggs available. At each new generation fresh host eggs were provided to these two modalities : 150 to 200 eggs for small carrying capacity (*modality S*) and 400 to 450 eggs for large carrying capacity (*modality L*). A third modality with lower resources was also tested under a slightly different protocol, see Supplementary Information S4.

Each modality of carrying capacity was replicated 20 times over 4 balanced time blocks, and populations were monitored during 10 generations.

#### Presence of positive density-dependent dispersal analysis

Using Approximate Bayesian Computation (ABC), we checked for the presence of positive density-dependent dispersal for *Trichogramma chilonis* in our experimental design by comparing two different scenarios, one with fixed dispersal and one with positive density-dependent dispersal (with a dependence of the form (7)). This analysis confirmed that the positive density-dependent dispersal scenario is the most likely, as suggested by a previous study with a comparable protocol (Morel-Journel et al., 2016). More details on the ABC analysis are presented as Supplementary Information S2.

#### Statistical analysis

Over the 40 experimental populations, 6 went extinct between generations 2 to 4 because of technical issues linked to climatic rooms where populations were reared. More extinctions occurred in the modality S (5, versus 1 in the modality L), which is consistent with theoretical predictions that small populations are more vulnerable to environmental stochasticity (Lande, 1993). Data corresponding to those populations were removed from the analyses. Each remaining population led to two propagation fronts (left and right), for which we calculated the number of colonized patches per population and generation. In total, 30 different fronts for modality S and 38 for modality L were analyzed.

To determine whether the carrying capacity influences propagation speed, we analyzed the number of colonized patches evolution for each front with a general linear model with mixed effects (GLMM, Bolker et al., 2009). The number of colonized patches was modeled with a Poisson law (log link). To account for the non-independence of data within each replicate across time and for environmental variance between blocks, we included replicate ID, front (left/right, nested with replicate) and block number as random effects on the slope. We tested two models corresponding to two hypotheses. The null model included only the generation as a fixed effect, thus considering that colonization speed was independent of carrying capacity. The alternative model included both generation and its interaction with experimental modality as fixed effects, accounting for a dependency between colonization speed and carrying capacity. We selected the best model based on Akaike Information Criterion (AIC): the best model has the lowest AIC. Then we computed the AIC weight *AIC*_*w*_ associated with the best model as follows, Δ(*AIC*) = *AIC* _*max*_ – *AIC*_*min*_ and 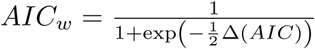.

## 3. Results

### Null model

As expected, whatever the probability of migration *p*_1_ (*p*_1_ = 0.25 or 0.5) the null IBM leads to results which are consistent with the Fisher-KPP reaction-diffusion model when *K* is not too small (here, *K* ≳ 20). Namely, the mean speed 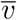 obtained with IBM simulations is almost independent of *K*. For smaller values of *K*, the behaviors of the two modeling frameworks tend to diverge due to the demographic stochasticity in the IBM. Whereas the propagation speed is always completely independent of *K* in the Fisher-KPP model, the simulations of the IBM lead to a positive and strongly increasing dependence between *K* and *v* for small values of *K*. These results are depicted in Fig. 4a.

**Figure 4:**
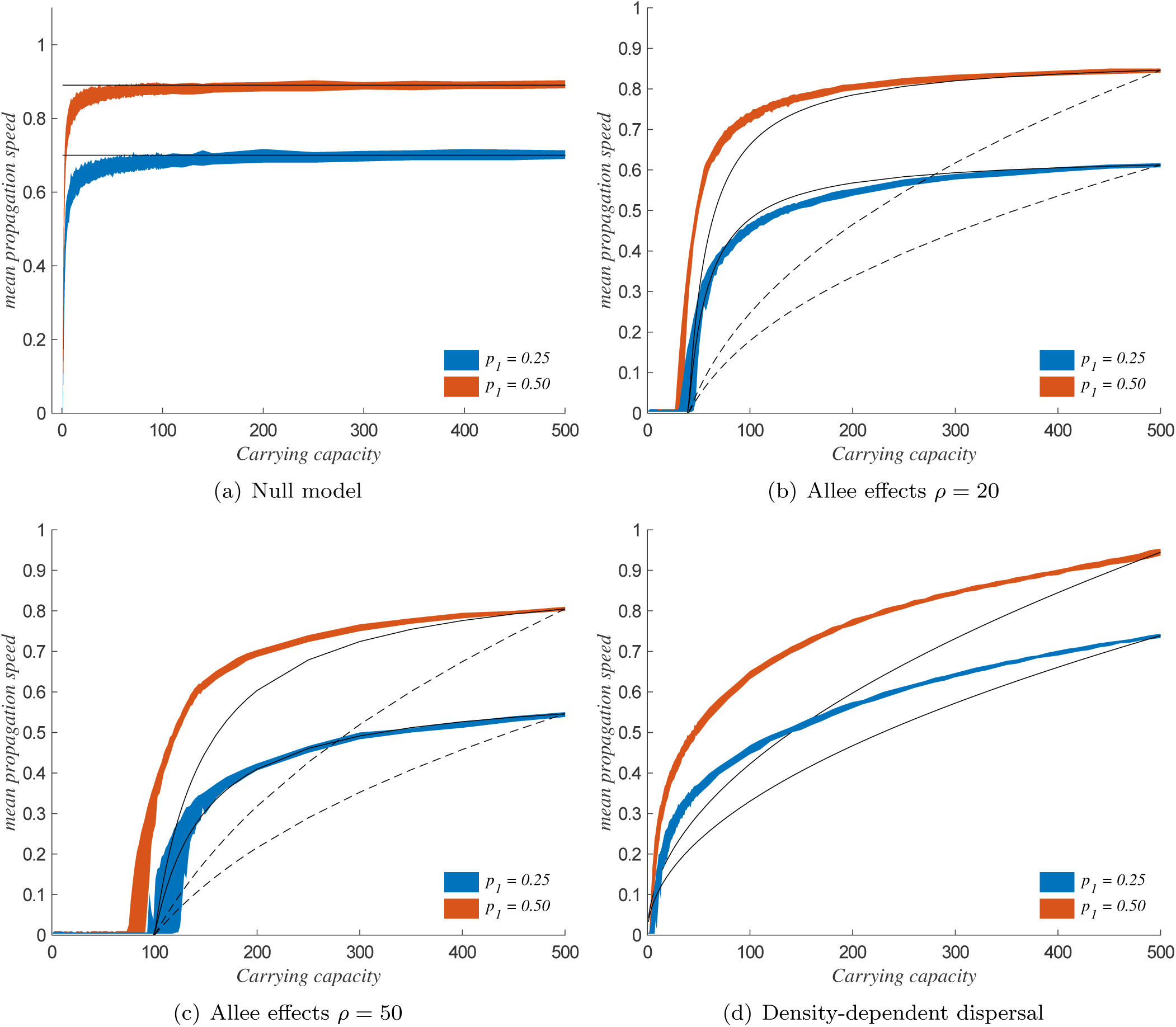
Propagation speed vs carrying capacity: IBM simulations and predictions of the reaction-diffusion theory. The shaded regions correspond to 99% confidence intervals for the mean speed obtained with individual-based simulations after 300 generations, with 200 replicate simulations for each scenario. The plain lines correspond to the propagation speed given by the analytical theory: formulas (2) for panel (a), formula (8) for panels (b) and (c) and formula (9) for panel (d). In panels (b),(c), the dashed lines correspond to the theoretical propagation speed obtained with classical growth function (3). In all cases, we set *R* = *e*′ in the IBM simulations and *r* = 1 in the reaction-diffusion framework. The diffusion coefficients *D* in the reaction-diffusion framework were fixed such that, in each scenario, the theoretical speed equals the average IBM speed for *K* = 500. The parameter values *p*_1_ in panel (d) (positive density-dependent dispersal) correspond to a population 𝒪_*i*_ = 500 individuals in formula (7).

### Strong Allee effect

As explained in Section 2.1, the standard form of growth function (3) usually used in reaction-diffusion models is not adapted to our study. However, it leads to a simple explicit formula for the propagation speed, which can be adapted to derive a formula for the propagation speed with the new form of growth function (4) that we proposed in Section 2.1. Namely, with the functional form (3), the propagation speed is 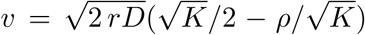, see Hadeler and Rothe (1975) for *K* = 1 and e.g., Keitt et al. (2001) and Roques (2013) for other values of *K*. Replacing *r* by 4 *r K/*(*K – ρ*)^2^ in this formula, we obtain a new formula for *v* for growth functions of the form (4):

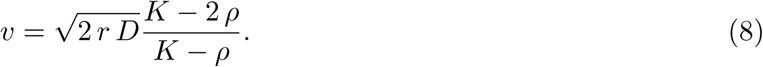

Thus, *v* is an increasing function of *K*, and converges to a finite value 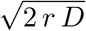 as *K → ∞*. Propagation occurs (*v >* 0) only when *K >* 2 *ρ*.

IBM simulations have been carried out with two values for the Allee threshold *ρ* (*ρ* = 20 and *ρ* = 50), and with two values for probability of migration (*p*_1_ = 0.25 or 0.5). The results are presented in Fig. 4b and 4c, together with the predictions of the reaction-diffusion approach. In agreement with the reaction-diffusion approach, propagation occurs only for values of *K ≳ ρ*, and after this threshold, the mean speed 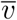 increases with *K*. We note that, in the IBM framework, propagation tends to occur for values of *K* slightly lower than predicted by the reaction-diffusion approach. The difference is more visible for large migration probability and Allee threshold (*p*_1_ = 0.5 and *ρ* = 50, see Fig. 4c). The stronger dependence between *K* and *v* is located around the threshold *K ≈* 2 *ρ*. Then, the curves converge towards horizontal asymptotes. The global shape of 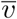 is close to that predicted by formula (8), but very different from that obtained with the standard formula corresponding to growth functions (3) (which predict that *v* increases like 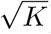, probably due to the dependence of the maximum per capita growth rate with respect to *K*, which does not occur in our IBM (see also Supplementary Information S1).

### Positive density-dependent dispersal

The reaction-diffusion framework proposed in Section 2.1, with a density-dependent diffusion coefficient *ϕ*(*u*) = *D u*, leads to a simple analytic formula for the propagation speed, which is obtained by adapting the formula in Newman (1980) (*K* = 1) to general values of *K*:

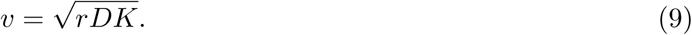

Thus, this model predicts that *v* increases like *K*^0.5^ when the diffusion coefficient is proportional to the population size. This is different from what was obtained with a strong Allee effect, where 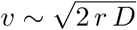 for large *K*.

In our IBM simulations, we fixed the constant *τ* to 1500 or 500 in (7) so that the probability of migration satisfies *p*_1_ = 0.25 or *p*_1_ = 0.5 when the population size equals 𝒪_*i*_ = 500, the maximum tested value for the carrying capacity. The results are presented in Fig. 4d. In all cases, we observe an increasing relationship between *K* and the mean speed 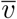. The dependence with respect to *K* is quite different from what was observed with a strong Allee effect. First, propagation always occurs, even for small values of *K*. Second, the mean speed 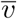 does not seem to converge towards an horizontal asymptote until it reaches its maximum possible value (1 in this framework where *δ*_*x*_ = *δ*_*t*_ = 1). A polynomial fit shows that 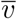 is approximately proportional to *K*^0.3^, which is qualitatively close to the prediction of the reaction-diffusion framework (sublinear increase), but quantitatively different, probably due to the differences in the assumptions on the form of the density-dependence.

Experimental results confirmed the positive relationship between carrying capacity and invasion speed, for a population displaying positive density-dependent dispersal but no Allee effect (Morel-Journel et al., 2016, see also Supplementary Information S2 for the ABC analysis of dispersal in *T. chilonis*). Fig. 5 (a) depicts the mean number of colonized patches over the 10 generations, for the two modalities of carrying capacity. Based on AIC selection, the best model estimates different slopes for the two modalities of carrying capacity (Δ(*AIC*) = 2.363, *AIC*_*w*_ = 0.765 ; interaction between generation and experimental modality: z value=-2.206, p-value=0.0274). In our experimental data, over the 10 generations, propagation fronts progressed with a speed of 0.17 patch per generation in the L modality against 0.13 patch per generation in the S modality; see Fig. 5 (b). From a quantitative viewpoint, when the *R* parameter in the simulation model is fixed so that the average propagation speed is 0.17 for the large modality (*K* = 400 in the IBM), we get a speed of 0.14 patch per generation for the small modality (*K* = 200 in the IBM). This is detailed in Supplementary Information S3.

**Figure 5:**
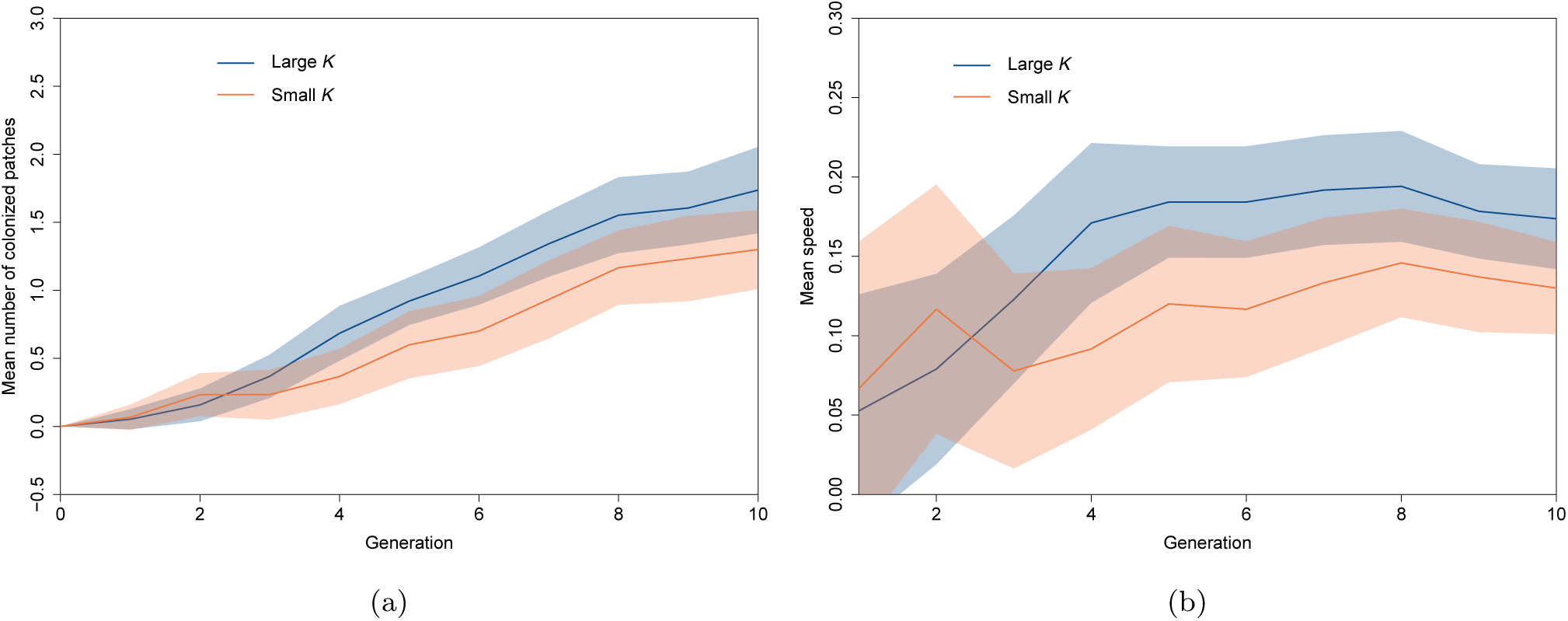
Mean number of colonized patches for the 68 replicate fronts over 10 generations (a); and corresponding mean speed (b). The red line corresponds to the small carrying capacity 150-200 host eggs, and the blue line corresponds to the large carrying capacity 400-450 host eggs. The red envelope is the 95% confident interval for the small modality and the blue envelope is 95% confidence interval for the large carrying capacity.

## 4 Discussion

### Different approaches, same conclusions

We used theoretical, simulation, and experimental approaches to investigate the influence of the carrying capacity *K* on the propagation speed of a population. These approaches revealed that the three biological mechanisms examined here could, under different conditions, lead to a positive relationship between *K* and *v*.

Positive density-dependent dispersal led to this increasing relationship under the three considered frameworks, independently of parameter values. However, the difference between the number of colonized patches for the S and L modalities in the experiment remained small. Theoretical and simulation results indicate that in the presence of positive density-dependent dispersal, the mean propagation speed is a concave (sublinear) function of *K* (Fig. 4d), with therefore a lower effect of *K* as *K* becomes large. We expect that lower values of *K* for the small modality would have led to significantly lower speeds. A posterior experiment that we conducted for another purpose and with a different protocol provides additional support to this conclusion (see Supplementary Information S4). Additional comparisons with simulation results (see Supplementary Information S3) also suggest that the difference between the S and L modalities should be more visible after more generations.

A Strong Allee effect led to similar conclusions to positive density-dependent dispersal under our theoretical and simulation frameworks but was not investigated experimentally. To disentangle the effect of *K* from the effect of the growth rate on the propagation speed, we proposed a new reaction-diffusion model for the strong Allee effects. With this model, contrarily to common approaches (Hadeler and Rothe, 1975; Lewis and Kareiva, 1993; Turchin, 1998; Keitt et al., 2001; Barton and Turelli, 2011), the maximal per capita growth rate is independent of *K*. With this new function, the shape of the propagation speed as a function of *K, v*(*K*), is closer to what we obtained with the simulation approach (Fig. 4b, and 4c); indeed by construction *K* does not impact the reproduction step in the IBM. In the presence of a strong Allee effect, our results indicate that the stronger dependence between *v* and *K* occurs when *K* is approximately two times larger than the Allee threshold *ρ*. This corresponds to the beginning of the propagation for the population.

In the absence of Allee effect and of density-dependent dispersal, theoretical and simulation approaches lead to consistent conclusions. When the carrying capacity is not too small the speed *v* is constant and independent of *K*. But, in the simulation approach, there is a strong relationship between the carrying capacity and the propagation speed for small populations. This dependency is induced by the stochasticity of the reproduction and dispersal step in the simulation approach. Such processes are expected to occur in all populations, but their impact is stronger in small populations (Gabriel and Bürger, 1992). This result highlights that even without any specific demographic mechanism, propagation speed may depend on the carrying capacity in small populations.

### Experimental studies with macro-organisms

Several recent studies compared the propagation properties of theoretical reaction-diffusion models *vs.* laboratory experiments. Yet they all used micro-organisms as biological models with large number of individuals (Giometto et al., 2014; Gandhi et al., 2016). These works have shown that the Fisher-KPP model is consistent with experimental speed when populations are not subject to any particular demographic process such as the Allee effect. Still like us they highlight the need for taking stochasticity into account in order to accurately predict speed, especially in small populations. For our experiment we chose to focus on a macro-organism with smaller population sizes (hundreds) experiencing important levels of demographic stochasticity. This allowed us to observe discrete numbers of individuals rather than densities of individuals.

These three distinct ecological mechanisms induced the same qualitative relationship – concave function – between carrying capacity and propagation speed. Interestingly, these mechanisms also share another common property: they make colonization more difficult when the source population is at low density, which impacts their propagation pattern. Therefore, the detection of a positive relationship between carrying capacity is not sufficient to discriminate between ecological mechanisms related to low-density dynamics, but is rather a qualitative indicator of a specific class of propagation dynamics.

### Carrying capacity impacting propagation speed, a proxy for pulled/pushed propagation?

Most of the time, studies on propagation dynamics focus on organisms with fast reproduction and dispersal. Range expansion in these types of populations is typically driven by individuals in small density at the edge of the range, which is described as pulled dynamics. In contrast, propagation dynamics are called pushed when they are influenced by the dynamics of individuals at intermediate or high density (Roques et al., 2012). Therefore, the dependency of the propagation speed on the carrying capacity should be an indicator of the pushed nature of a wave. Populations subject to strong Allee effects are known to experience pushed dynamics (Roques et al., 2012; Gandhi et al., 2016). Stochasticity was also described by Panja (2004) as leading to “weakly pushed” dynamics (i.e., pushed dynamics that would converge to pulled dynamics in the limit of infinite population sizes). Based on the results of our study, we argue that positive density-dependent dispersal also leads to pushed propagation patterns. The pushed nature of the dynamics implies that the individuals at the edge of the populations are not able to fully drive the propagation. This may be because their colonization potential at low density is not efficient, e.g. Allee effects impacting reproduction or the stochasticity impacting reproduction, mortality and dispersal. For positive density-dependent dispersal individuals at the edge of the front do not produce enough dispersers to allow further propagation, thus causing a pushed expansion pattern. Conversely, independence of *v* with respect to *K* does not necessarily imply that the waves are pulled, but strongly suggests it.

Understanding the pulled or pushed nature of propagation is an important issue for forecasting spread and elaborating conservation or eradication strategies. Control or eradication strategies for populations subjected to an Allee effect have been well theorized. They are based either on the increase of the Allee threshold, or on reducing population size under the existing threshold through culling or biological control (Taylor and Hastings, 2005; Tobin et al., 2011). Unlike pulled expansions, pushed populations might also be managed indirectly, by altering the carrying capacity of the landscape. For instance, the “range pinning” in which expansion stops even in the presence of suitable habitat has been theoretically described for populations subject to a strong Allee effect (Keitt et al., 2001), and might be a general property of pushed expansion fronts. Recent theoretical work has also shown that pushed expansions can be halted by creating a barrier of unfavorable habitat even if this barrier has holes (Roques et al., 2008; Tanaka et al., 2017). In addition to these theoretical predictions, recent experimental results obtained in periodic environments show that expansion can be stopped by a succession of low-K habitats in the presence of density-dependent dispersal (Morel-Journel et al., 2018).

Feedbacks from re-introduction programs for conservation purposes also provide some valuable insight about expansion dynamics at low density. In many cases, populations that have been re-introduced failed to expand (or expanded very slowly), despite a positive demography, e.g., raptors in Great Britain (Carter et al., 2003; Mackrill et al., 2013) or large predators in North America (Hayward and Somers, 2009; Hornocker and Negri, 2009). This phenomenon was linked to the presence of an Allee effect (Hurford et al., 2006), to high philopatry (Mackrill, 2017), which often relates to density-dependent dispersal, or to poor dispersal abilities (Hornocker and Negri, 2009), which increases dispersal stochasticity. All these characteristics were likely to induce pushed propagation patterns. In such cases, the counter-intuitive measure of improving the local habitat to further increase the size of the source population might be an efficient way to promote spatial expansion. However, highly endangered populations subject to expensive re-introduction programs have often also suffered from massive habitat loss (Kramer-Schadt et al., 2005), so the remaining available habitat may not be sufficient to allow for pushed expansions to occur even if viable populations can be locally sustained. In this case, further re-introductions would be needed to establish enough population cores to ensure long-term persistence. In a second step, dispersal fluxes should be monitored to verify whether the population cores manage to achieve metapopulation dynamics, or whether the regular translocation of individuals should be maintained in the long term (Armstrong and Seddon, 2008).

This work is the first theoretical and empirical demonstration of the influence of carrying capacity on propagation speed. Our results suggest that this relationship is a common property of pushed waves, which are characterized by density-dependent colonization success. This finding raises innovative perspectives for the use of landscape properties for the management of pushed populations and for the optimization of re-introduction programs.

## 5 Acknowledgements

The authors are indebted to Samuel Soubeyrand for his help with ABC analyses and to Vincent Calcagno for significantly improving the manuscript. We thanks Matthieu Barbier for reviewing and recommending the manuscript, Yuval Zelnik and one anonymous reviewer provided insightful comments on the manuscript. MH was supported by a PhD fellowship funded by INRA, division Plant Health and Environment, and the Council of Provence Alpes Côte d’Azur region. The research leading to these results has received funding from the French Agence Nationale de la Recherche within the project NONLOCAL (ANR-14-CE25-0013) and the project TriPTIC (ANR-14-CE18-0002). This work was also funded by INRA grant “MEDIA”.

## 6 Statement of authorship

TMJ, EL, EV, LM, LR designed research. All authors contributed to design the experiment. MH and TMJ conducted the experiment and collected data. All authors contributed to analyze the data. MH and LR contributed to the mathematical analysis. MH, EV, LM, LR wrote the first draft. All authors contributed to revisions of the manuscript.

## 8 Data Accessibility statement

Data available from Zenodo repository: https://doi.org/10.5281/zenodo.1420074

